# Sympathetic blockage attenuates fasting-induced hepatic steatosis

**DOI:** 10.1101/2022.02.13.480240

**Authors:** Lin Zheng, Lijun Gong, Siyu Lu, Yiming Zhou, Zhigui Duan, Fang Wei, Guolin Li

## Abstract

**Background and Aim:** Although it is clear that the central nervous system coordinates whole-body metabolism, the neural mechanism for hepatic steatosis remains unclear. This study is aimed to explore the neural mechanism of fasting-induced hepatic steatosis.

**Methods:** Mice were pretreated with 6-hydroxydopamine to block sympathetic nerve activity before fasting, and explored the potential effects of chemical sympathectomy on fasting-induced hepatic steatosis and transcriptional changes.

**Results:** Prolonged fasting led to obvious hepatic steatosis, low core temperature, and similar effects to cold-induced white adipose lipolysis. The alterations in hepatic mRNA expression revealed that the hepatic lipid accumulation did not result from the increase of hepatic lipogenesis or the decrease of fatty acid oxidation but from the enhanced fatty acid uptake as indicated by the upregulation of CD36. Blockage of the sympathetic nervous system via chemical sympathectomy attenuated fasting-induced hepatic steatosis and suppressed CD36 upregulation in the liver, but did not obviously alter the expression of genes associated with lipogenesis or fatty acid oxidation.

**Conclusions:** These findings indicate that the sympathetic nervous system orchestrates the mechanism for fasting-induced hepatic steatosis via modulating CD36 expression and adipose fat trafficking into the liver, which provides clues to reveal new targets for fatty liver diseases.

**HIGHLIGHTS:**

- Prolonged fasting causes obvious hepatic steatosis
- Sympathectomy attenuates fasting-induced hepatic steatosis
- Sympathectomy attenuates steatosis via suppressing CD36 upregulation

**IN BRIEF:** Prolonged fasting contributes to sympathetic hyperactivity, but its role in the pathogenesis of hepatic steatosis remains unclear. Here, we found prolonged fasting led to obvious hepatic steatosis, low core temperature, and subsequent sympathetic stimulation that promoted white adipose lipolysis, hepatic upregulation of CD36, and adipose fat trafficking into the liver.

## INTRODUCTION

Nonalcoholic fatty liver disease (NAFLD) is emerging as a leading global health problem that affects approximately 25% of the world population(Younossi et al., 2019). The disease begins with the aberrant triglyceride (TG) accumulation in the liver, which can progress to steatohepatitis, advanced fibrosis, cirrhosis, and hepatocellular carcinoma(Haas et al., 2016; Loomba et al., 2021). In general, hepatic steatosis is stemmed from maladaptations of fatty acid synthesis, fatty acid oxidation, fatty acid uptake, or export of lipids from the liver(Cohen et al., 2011; Sozio et al., 2010). However, the molecular mechanism responsible for hepatic steatosis and its progression remains to be fully elucidated.

Owing to the regulatory network of lipid metabolism being complex, multiple factors may contribute to the pathogenesis of NAFLD(Reddy and Rao, 2006; Sozio et al., 2010). It is now clear that a large number of dietary, chemical, and genetic factors can elicit hepatic steatosis and its progression(Van Herck et al., 2017; Zhong et al., 2020). Given that the central nervous system coordinates whole body metabolism(Bruinstroop et al., 2014; Myers and Olson, 2012; Yi et al., 2010), neuromodulation has recently emerged in the pathogenesis of NAFLD(Adori et al., 2021; Loomba et al., 2010; Miyamoto et al., 2012; Oben et al., 2010). Growing evidence demonstrated that the autonomic nervous system can either directly regulate hepatic lipid accumulation(Amir et al., 2020; Bruinstroop et al., 2014; Edinburgh and Frampton, 2020; Guarino et al., 2017; Yi et al., 2010) or indirectly modulate hepatic steatosis through governing the release of adipokines(Miyamoto et al., 2012; Oben et al., 2010; Wang et al., 2020; Warne et al., 2011; Zeng et al., 2015). Overexpression of β-adrenergic receptors in cultured hepatocytes increases cellular lipid accumulation, while β-adrenergic blockers prevent the development of hepatic steatosis(Ghosh et al., 2012). Diet-induced obesity mice characterize hepatic steatosis and high activity of sympathetic nerves, while blockage of sympathetic nerves abolishes obesity-induced steatosis(Hurr et al., 2019). Intriguingly, prolonged fasting has been reported to cause hepatic steatosis(Li et al., 2018) and elevate sympathetic nerve activity (Cohen et al., 2011; Knehans and Romsos, 1983). However, whether the elevated sympathetic nerve activity contributes to hepatic lipid accumulation or just a co-consequence of hepatic steatosis induced by fasting remains unclear.

In this study, we pretreated mice with 6-hydroxydopamine (6-OHDA) to block sympathetic nerve activity before fasting, and explored the potential effects of chemical sympathectomy on fasting-induced hepatic steatosis and transcriptional regulation. Our results indicate that the sympathetic nervous system (SNS) orchestrates the mechanism for fasting-induced hepatic steatosis through modulating the expression of hepatic lipid transporter cluster of differentiation 36 (*Cd36*) and adipose fat trafficking into the liver, which provides clues to reveal new targets for fatty liver diseases.

## METHODS

### Ethics Statement

The human study was approved by the Ethics Committee of the Hunan Normal University. All study volunteers signed the informed consent forms before their inclusion in the project.

### Animals

All mouse studies were approved by the Animal Care and Use Committee of Hunan Normal University and performed following the Laboratory Animal Resources guidelines. Six-week-old male C57BL/6 mice and standard rodent chow diets were purchased from Hunan SJA Laboratory Animal Co., Ltd. Mice were housed at 22–25°C on a 12-hour light/12-hour dark cycle with ad libitum access to food and water. All experiments were started with 8-week-old mice.

### Fasting models

Mice were randomly grouped to control (Con) group, acute fasting for 24 hours (fasting) group, 6-OHDA treatment group, 6-OHDA+fasting treatment group. All mice were singly housed for 2 weeks before study initiation to allow for acclimation to the animal facility. The Control group mice were allowed unrestricted access to food. The mice of fasting group underwent acute fasting for 24 hours before being sacrificed. The mice of 6-OHDA group were given intraperitoneal injection of 6-OHDA (100 mg/kg b.w.) 24 hours before being sacrificed. In 6-OHDA+fasting group, the mice were intraperitoneal injected with 6-OHDA, followed by an acute fasting for 24 hours before being sacrificed.

### Blood Glucose and Core Temperature Tests

Mice circadian blood glucose was measured with a Glucometer (Cofoe, Changsha, Hunan) by tail bleeds. The core body temperature was measured rectally with a Thermometer equipped with a RET-3 mouse rectal temperature probe (WPI, Sarasota, FL).

### Serum Biochemical Assays

Serum triglyceride (TG) and non-esterified fatty acids (NEFA) were assessed by commercial assay kits (Jiancheng, Nanjing, China) and monitored with a NanoDrop 2000 Spectrophotometer (Thermo Scientific, Wilmington, DE).

### Liver Lipid Assay

For analysis of liver TG and NEFA content, 100 mg of the frozen liver was homogenized in 900 μl of RIPA lysis buffer + 1% Triton X-100. After centrifugation, the supernatants were diluted 10 folds. Then, lipids were quantified using commercial assay kits (Jiancheng, Nanjing, China) and the concentrations of protein were determined using a commercial BCA protein assay kit (Servicebio, Wuhan, China).

### Total RNA Extraction, cDNA Synthesis, and Real-time PCR

Total RNA was extracted from frozen tissues using TRIzol reagent according to the manufacturer’s instructions. The purities and concentrations of total RNA samples were determined by NanoDrop 2000 Spectrophotometer (Thermo Scientific, Wilmington, DE). 200 ng of total RNA was reverse transcribed using cDNA Synthesis Mix (Takara, Dalian, China). Real-time PCR was carried out in an Applied Biosystems QuantStudio 5 Real-Time PCR System (ABI, Warrington, UK) with SYBR Green PCR master mix (Takara, Dalian, China) and gene-specific primers. The sequences for the forward and reverse primers used to quantify mRNA are listed in Supplementary Table 1. The following conditions were used for real-time PCR: 95°C for 10 min, then 95°C for 15 sec, and 60°C or 1 min in 40 cycles. The 2^−ΔΔCT^ method (Pfaffl, 2001) was used to analyze the relative changes in gene expression normalized against *Gapdh* expression. qRT-PCR experiments were designed and performed according to Minimum Information for Publication of Quantitative Real-Time PCR Experiments (MIQE) guidelines (Bustin et al., 2009).

### Western Blot Analysis

Frozen tissues were lysed in RIPA buffer supplemented with Halt Protease and Phosphatase Inhibitor Cocktail (Servicebio, Wuhan, China) and 1 mM PMSF. Protein concentrations were determined using BCA Protein Assay Kit (Servicebio, Wuhan, China) by a microplate reader (PERL ONG, Beijing, China). Twenty micrograms of protein per lane were loaded onto an 8-12% polyacrylamide Gel and then transferred to PVDF membranes using an electrophoresis chamber (Millipore, Shanghai, China) for 1.5 h. Membranes were blocked in 5% non-fat milk Tris-buffered saline (TBS) followed by overnight incubation in the primary antibody at 4°C. Primary antibodies used are as follows: rabbit PKA antibody (#3927S, CST, Danvers, MA, USA), Rabbit phosphorylated PKA antibody (#PA5-1055515, Invitrogen, Carlsbad, California, USA), rabbit HSL antibody (#41075, CST, Danvers, MA, USA), and rabbit Phosphorylated HSL antibody (#PA5-105769, Invitrogen, Carlsbad, California, USA). Following primary antibody incubation, blots were washed four times for 10 min each time with TBS and 0.1% Tween 20 and incubated with an anti-rabbit IgG HRP-conjugated secondary antibody (#AS014, ABclonal, Wuhan, China). Blots were stripped with Enhanced Chemiluminescent Liquid (ECL, NCM, Suzhou, China) and re-probed with anti-Tubulin for normalization (#AF7011, Affinity Biosciences, Suzhou, China). Blot imaging was performed on a chemiluminescence analyzer (Tanon-5200 Multi, Shanghai, China).

### Hematoxylin and Eosin (H&E) Staining

All tissues were fixed in 4% paraformaldehyde for 24 h at room temperature, dehydrated, and embedded into paraffin. The tissues were sectioned into 2 μm slices and stained with hematoxylin and eosin (H&E). Digital images were collected at 40× magnification with a Niko Eclipse E100 upright optical microscope (Nikon, Japan) equipped with a microscopic examination, and NIKON DS-U3 imaging system (Nikon, Japan). The images shown are representative results of at least three biological replicates.

### Statistical Analysis

All results are expressed as means ± SEM. Significance was determined by *t*-test or one-way ANOVA with Bonferroni posttest using Prism 7.0 software (GraphPad Software,). A *P* value of less than 0.05 was considered significant.

## RESULTS

### Fasting contributes to hepatic steatosis and white adipose hydrolysis

Consistent with present evidence that prolonged fasting leads to lipid accumulation in the liver (Li et al., 2018), our results illustrated that fasting for 24 hours in mice caused obvious hepatic steatosis (Figure 1A and 1B). The fasting also increased the serum levels of TG and NEFA (Figure 1C and 1D) and hepatic NEFA (Figure 1E). By contrast, the blood glucose decreased gradually with the extension of fasting time (Figure 1F). Given that fasting in mice leads to the primary fuel sources shift from carbohydrates to lipids (Jensen et al., 2013), these results are logical. Intriguingly, we noticed that fasting lowered the mouse rectal temperature, promoted the phosphorylation of hormone-sensitive lipase (HSL), while declining the weight ratio of white adipose to the liver (Figure 1G-1J), indicating a seemed low temperature-induced white adipose lipolysis as repeatedly reported in cold-induced adipose browning and lipolysis (Champigny and Ricquier, 1990; Knehans and Romsos, 1983; Landsberg et al., 1984; Lowell and Spiegelman, 2000; Puigserver et al., 1998). Besides, the declined weight ratio of white adipose to the liver may suggest that fasting may promote fat mobilization and adipose fat trafficking into the liver.

**Figure 1.**
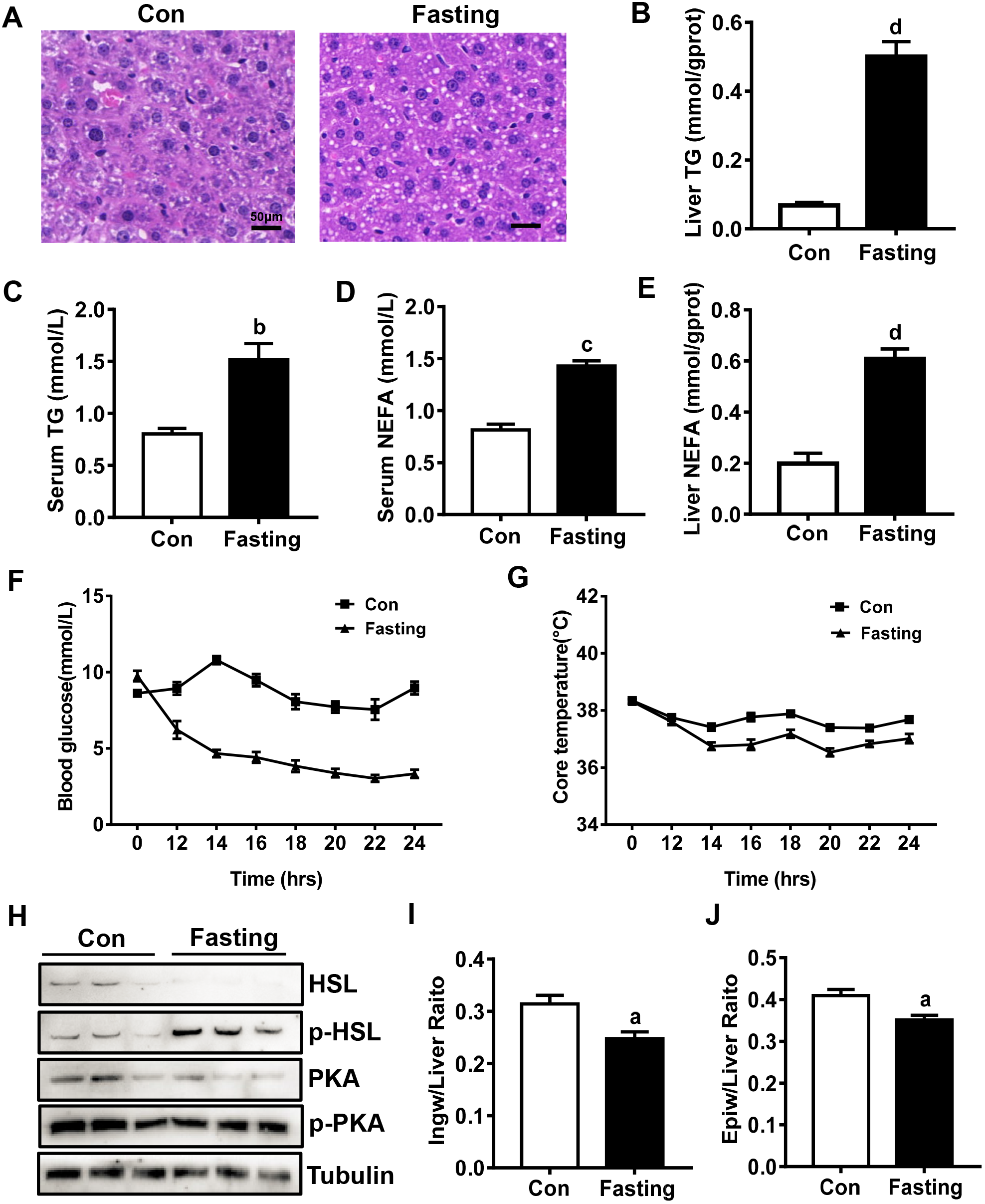
Fasting contributes to hepatic steatosis and adipose hydrolysis. A) Representative H&E staining of liver sections from control (Con) or acute fasting for 24 h (fasting) mice. Scale bar: 50 μm. B-G) The effects of acute fasting on the levels of liver TG (B), serum TG (C), serum NEFA (D), liver NEFA (E), blood glucose (F), and rectal temperature (G) in mice. (n=5) H) Liver extracts were immunoblotted with the indicated antibodies. I-J) The weight ratio of IngW (I) and EpiW (J) to the liver in mice treated with or without acute fasting. (n=5) Data are presented as mean ± SEM. Different lowercase letters indicate statistical significance in fasting versus Con (a, *P* < 0.05; b, *P* < 0.01; c, *P* < 0.005; and d, *P* < 0.001). Con, control; EpiW, epididymal white adipose tissue; HSL, hormone-sensitive lipase; IngW, inguinal white adipose tissue; NEFA: non-esterified fatty acids; pHSL, phosphorylated hormone-sensitive lipase; PKA, protein kinase A; pPKA, phosphorylated protein kinase A; TG, triglyceride.

### Fasting altered the expression of mRNA associated with hepatic lipid metabolism homeostasis

As we know, hepatic steatosis arises from an imbalance between TG formation and removal(Cohen et al., 2011; Sozio et al., 2010), and fatty acids from diet, de novo synthesis, and adipose tissue are three sources for hepatic TG formation(Cohen et al., 2011). Therefore, to explore potential mechanisms orchestrated fasting-induced hepatic steatosis, we analyzed the mRNA expression of genes responsible for lipogenesis, lipid transport, and fatty acid oxidation in the liver and primary hepatocytes. The results displayed that fasting in mice significantly decreased the expression of hepatic lipogenic genes (Figure 2A), while increased these of fatty acid oxidation and lipid transport (Figure 2B and 2C). These data are consistent with the above results and suggest that white adipose lipolysis rather than lipid de novo synthesis or the suppression of fatty acid oxidation contributes to fasting-induced hepatic steatosis.

**Figure 2.**
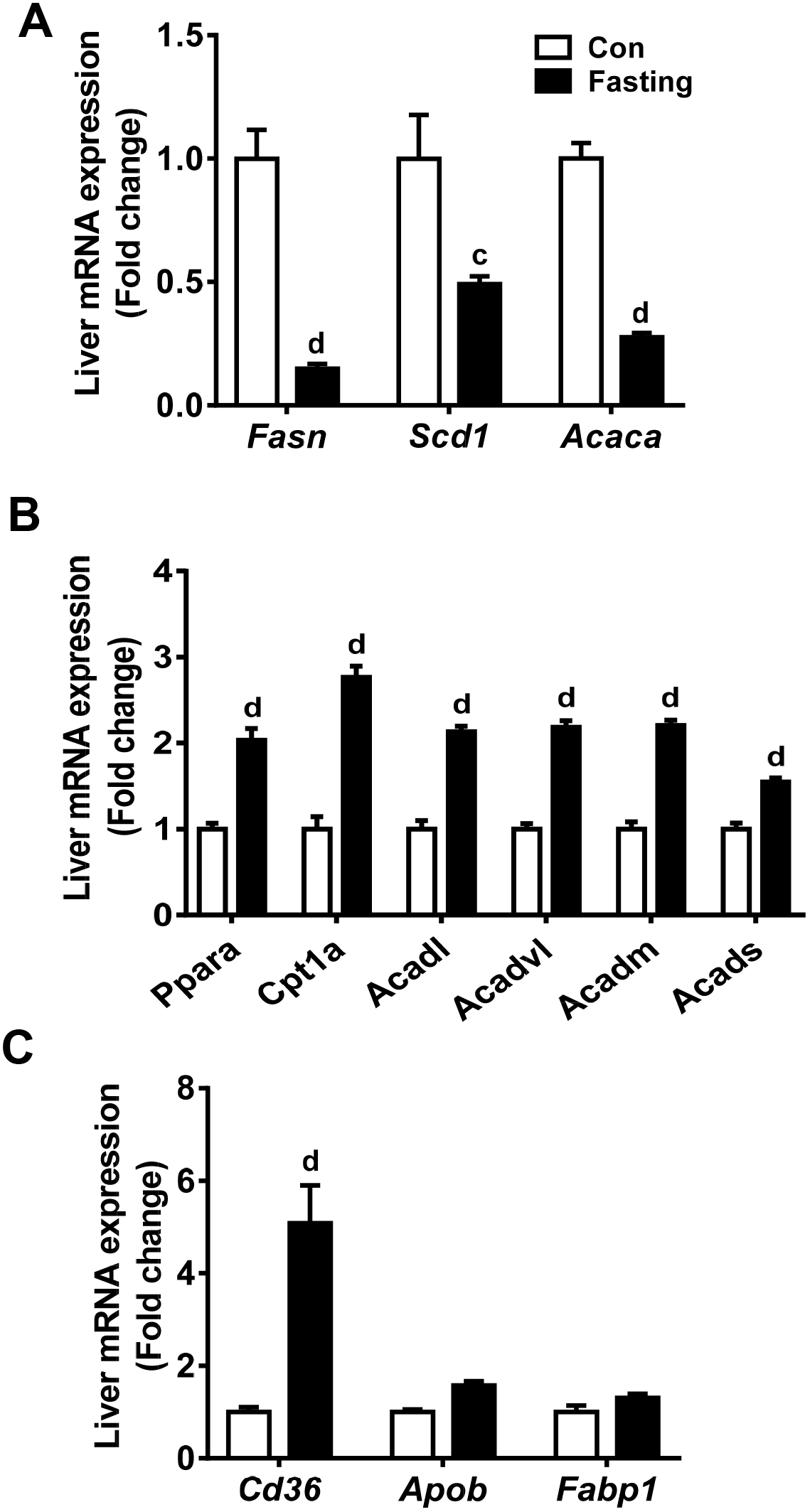
Fasting alters hepatic lipid metabolism homeostasis. A-C) The mRNA expression (fold change to control mice) of genes responsible for lipogenesis (A), fatty acid oxidation (B), and fatty acid transport (C) in the liver of mice treated with or without acute fasting. (n=5) Data are presented as mean ± SEM. Different lowercase letters indicate statistical significance in fasting versus Con (a, *P* < 0.05; b, *P* < 0.01; c, *P* < 0.005; and d, *P* < 0.001). Con, control.

### Sympathectomy attenuates fasting-induced hepatic steatosis

Since fasted mice displayed a low temperature-induced adipose lipolysis, and the stimulation of the SNS is the canonical mechanism for cold-induced adipose browning and lipolysis (Landsberg et al., 1984; Lowell and Spiegelman, 2000), we suspected that fasting-induced steatosis in mice might be associated with the SNS stimulation. Therefore, we further estimated the effect of fasting after the elimination of sympathetic nerve activity by chemical sympathectomy with 6-OHDA (Thoenen and Tranzer, 1968). As expected, the sympathectomy attenuated fasting-induced hepatic steatosis (Figure 3A-3C). Although it did not obviously alter serum levels of TG and NEFA (Figure 3D and 3E), the fasting-induced low temperature and blood glucose improved (Figure 3F and 3G), and the decrement ratio of white adipose to the liver was inhibited (Figure 3H and 3I). These data suggest that the sympathectomy ameliorates fasting-induced steatosis through the suppression of fat mobilization and adipose fat trafficking into the liver.

**Figure 3.**
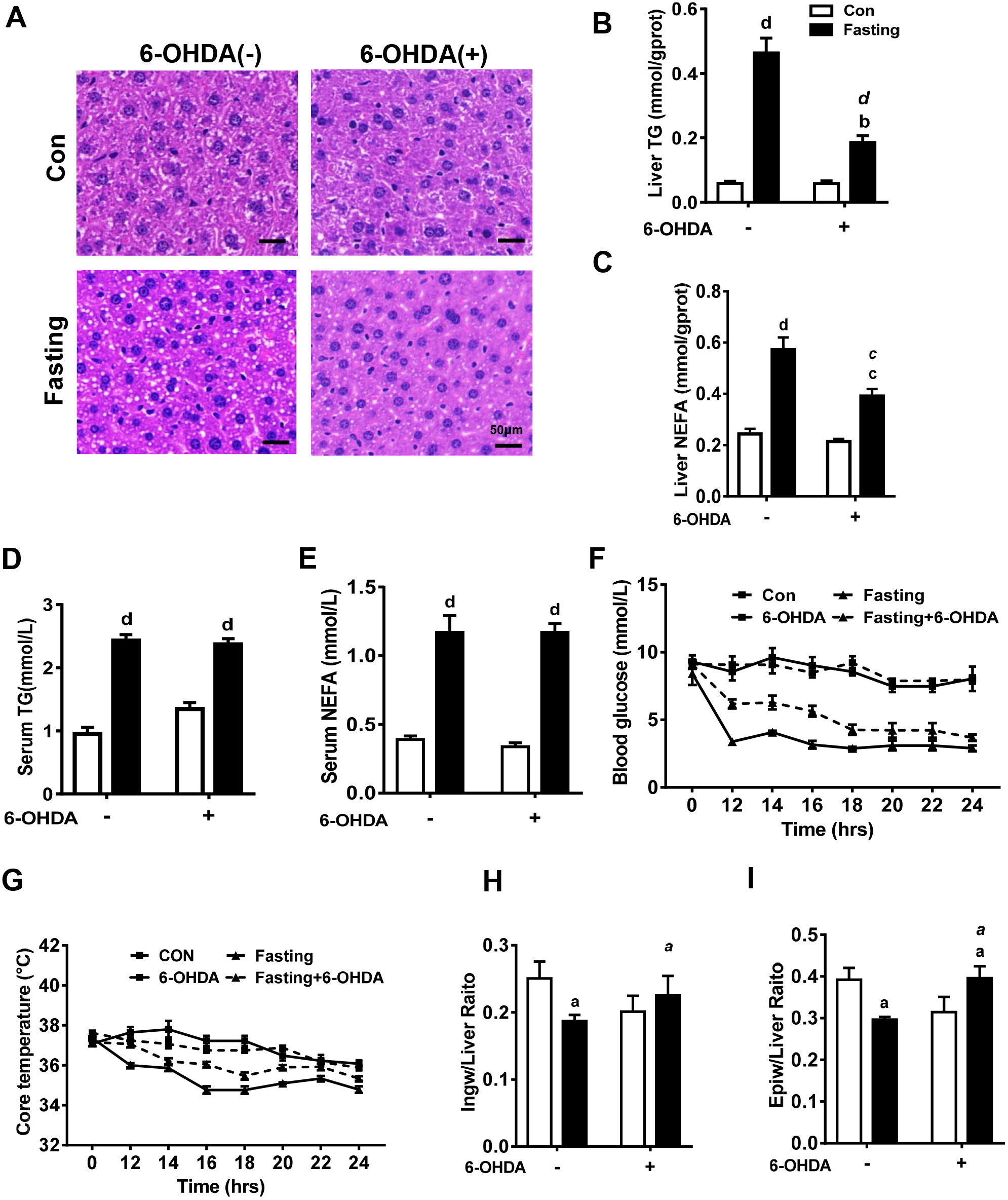
Sympathectomy attenuates fasting-induced hepatic steatosis. A) Representative H&E staining of liver sections from mice treated with 6-OHDA and/or fasting. Scale bar: 50 μm. B-G) The effects of acute fasting on the levels of liver TG (B), liver NEFA (C), serum TG (D), serum NEFA (E), blood glucose (F), and rectal temperature (G) in mice with or without 6-OHDA treatment. (n=5) H-I) The effects of acute fasting on the weight ratio of IngW (H) and EpiW (I) to the liver in mice treated with or without 6-OHDA. (n=5) Data are presented as mean ± SEM. Different lowercase letters indicate statistical significance (a, *P* < 0.05; b, *P* < 0.01; c, *P* < 0.005; and d, *P* < 0.001). Normal letters show the effects of fasting (Fasting versus Con within the same 6-OHDA treatment), and *italic* letters show the effects of 6-OHDA [6-OHDA(+) versus 6-OHDA(−) within the same feeding treatment]. Con, control; EpiW, epididymal white adipose tissue; IngW, inguinal white adipose tissue; NEFA: non-esterified fatty acids; 6-OHDA, 6-hydroxydopamine; TG, triglyceride.

### Sympathectomy attenuates fasting-induced upregulation of mRNA associated with lipid transport

To confirm the above concept, we also analyzed the mRNA expression of genes responsible for lipogenesis, lipid transport, and fatty acid oxidation in the liver and primary hepatocytes. The results illustrated that blockage of the SNS did not obviously change the process of lipogenesis and fatty acid oxidation, but suppressed fasting-induced upregulation of *Cd36* and other genes associated with fatty acid transport in mice (Figure 4A-4C). Therefore, these results reinforced the concept that adipose fat trafficking into the liver during prolonged fasting might be the primary mechanism for fasting-induced steatosis.

**Figure 4.**
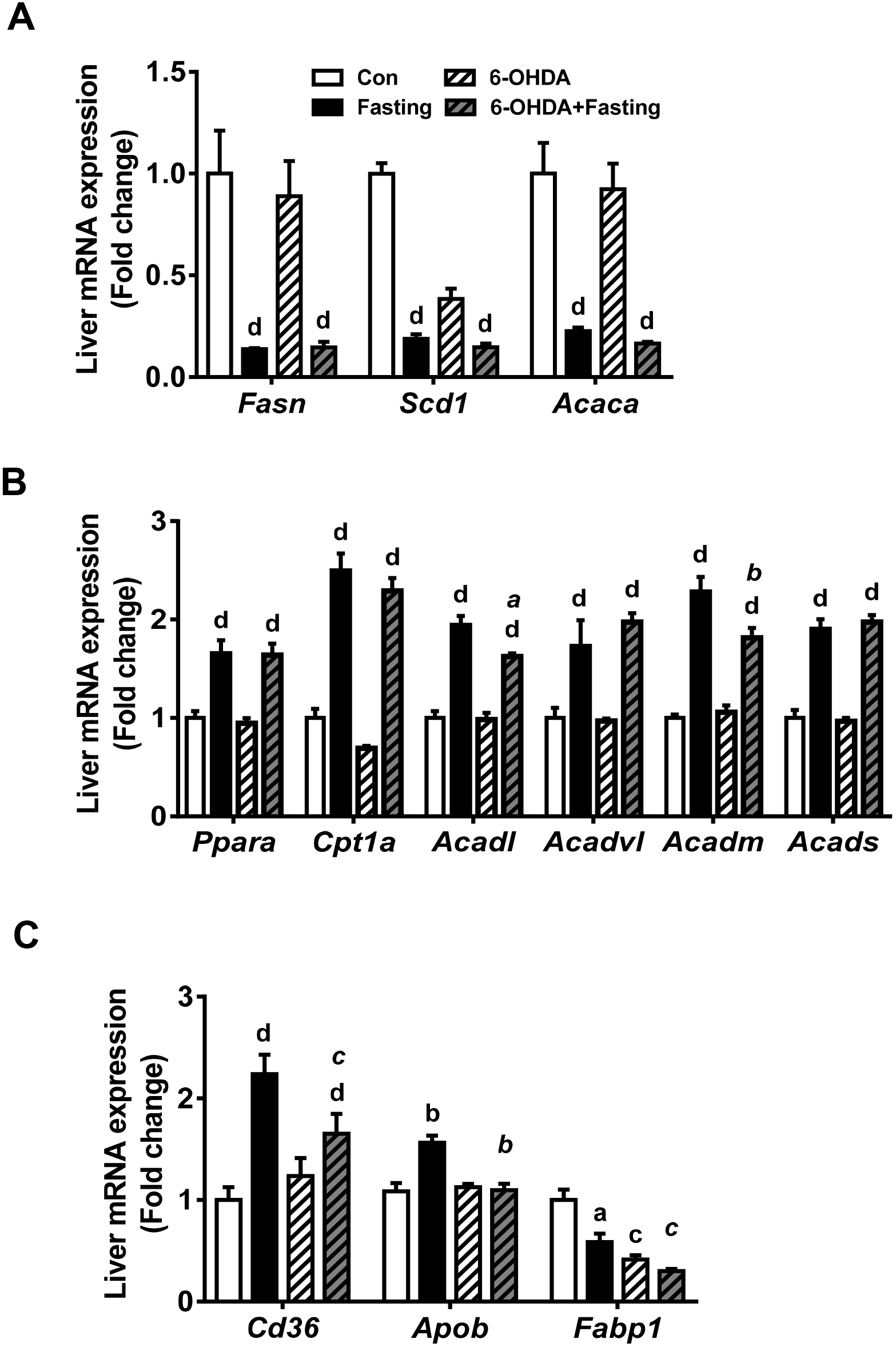
Sympathectomy attenuates the upregulation of fatty acid transporters induced by fasting. A-C) The effects of 6-OHDA on mRNA expression (fold change to control mice without 6-OHDA treatment) of genes responsible for lipogenesis (A), fatty acid oxidation (B), and fatty acid transport (C) in the liver of mice treated with or without acute fasting. (n=5) Data are presented as mean ± SEM. Different lowercase letters indicate statistical significance in fasting versus Con (a, *P* < 0.05; b, *P* < 0.01; c, *P* < 0.005; and d, *P* < 0.001). Con, control; 6-OHDA, 6-hydroxydopamine.

## DISCUSSIONS

The primary finding from this study is that the SNS orchestrates fasting-induced hepatic steatosis. Although the concept that the SNS stimulation in stress conditions is present to prepare the animal for “fight or flight” by mobilizing fatty acids from adipose tissue has been revealed by Cannon in 1915(Cannon, 1915), and prolonged fasting can increase the plasma levels of epinephrine or norepinephrine and white adipose lipolysis has been repeatedly reported(Cohen et al., 2011; Knehans and Romsos, 1983; Rayner, 2001; Vollmer and Skott, 2002), there is no evidence directly link the SNS to fasting-induced hepatic steatosis. In this study, we found sympathectomy attenuated fasting-induced steatosis, which provided the first evidence that SNS orchestrates fasting-induced hepatic steatosis. This finding is consistent with previous studies that the sympathetic stimulation by aging, obesity, or genetic modulation contributed to hepatic steatosis(Ghosh et al., 2012; Hurr et al., 2019) and reinforces the concept that the SNS regulates hepatic lipid homeostasis(Amir et al., 2020; Bruinstroop et al., 2014; Edinburgh and Frampton, 2020; Guarino et al., 2017; Yi et al., 2010).

Mechanistically, the suppression of CD36 expression may mediate sympathectomy-induced amelioration of steatosis in fasted mice. In this study, we found that fasting evidently upregulated the expression of *Cd36* and promoted hepatic lipid accumulation, while blockage of the SNS suppressed the *Cd36* upregulation and steatosis. Consistent with these results, a recent study in diet-induced obesity mice demonstrated that CD36 expression was modulated by hepatic sympathetic innervation, while hepatic sympathetic denervation suppressed CD36 expression and fatty liver(Hurr et al., 2019). As we know, CD36 is a fatty acid transporter in hepatocyte membrane(Zhou et al., 2008), and the upregulation of CD36 directly contributes to the development of fatty liver under conditions of elevated free fatty acids by modulating the rate of fatty acid uptake(Wilson et al., 2016). Therefore, CD36 is considered a key driver for fatty liver and regulator of insulin sensitivity(Rada et al., 2020; Wilson et al., 2016; Zhou et al., 2008). Intriguingly, among the four primary factors directly regulating the balance of lipid acquisition and removal(Cohen et al., 2011; Sozio et al., 2010), we found that sympathectomy only obviously altered fatty acid uptake by downregulating the gene expression of fatty acid transporters including CD36. In this context, it is logical to suggest that the mechanism of sympathetic blockage in improving hepatic steatosis may be derived from the suppression of CD36 expression.

In this fasting-induced steatosis mouse model, the expression pattern of genes involved in lipogenesis and fatty acid oxidation favors the removal rather than accumulation of lipid, which seems contradict to the pathogenesis of fatty liver(Cohen et al., 2011; Sozio et al., 2010). However, if we consider the decreased lipogenesis and increased fatty acid oxidation as a compensatory adaptation to hepatic excessive uptake of fatty acid from the circulation, it is logical. In fasted mice, we did find elevated NEFA levels in the liver, which would undoubtedly inhibit de novo synthesis of fatty acid. According to previous studies, under fasted state, the nuclear receptor peroxisome proliferator-activated receptor alpha (PPARα) is activated by fatty acids, which subsequently promote fatty acid oxidation (Evans et al., 2004; Gottlicher et al., 1992; Keller et al., 1993).

In summary, the present work uncovered novel pathogenesis of fasting-induced hepatic steatosis. Acute fasting was shown to rapidly develop hepatic steatosis, probably by resulting in hypothermia and subsequent SNS stimulation, which promoted white adipose lipolysis, hepatic upregulation of CD36, and adipose fat trafficking into the liver. This novel pathogenesis opens a new window for revealing new mechanisms of NAFLD and would hold promise for discovering new therapeutic targets for NAFLD.

## ACKNOWLEDGMENTS

We thank Li Liu, Yu Liang, and Can Zhou for their assistance with tissue collection. G.L. was supported by the National Natural Science Funds of China (31871198), and the Opening Fund of The National & Local Joint Engineering Laboratory of Animal Peptide Drug Development (Hunan Normal University, National Development and Reform Commission). F.W. was supported by the National Natural Science Funds of China (81903138). The funding sponsors had no role in the writing of the manuscript, and in the decision to submit the manuscript for publication.

## AUTHOR CONTRIBUTIONS

G.L. designed and supervised the experiments. L.Z., L.G., S.L., Y.Z., Z.D., and F.W. conducted experiments. L.Z., F.W., and G.L. performed the data analysis. G.L. and F.W. interpreted results. G.L. wrote the manuscript. G.L., F.W., and L.Z. revised the manuscript.

## DECLARATION OF INTERESTS

The authors declare no competing interests.

## FIGURE LEGENDS

**Supplementary Table 1.**
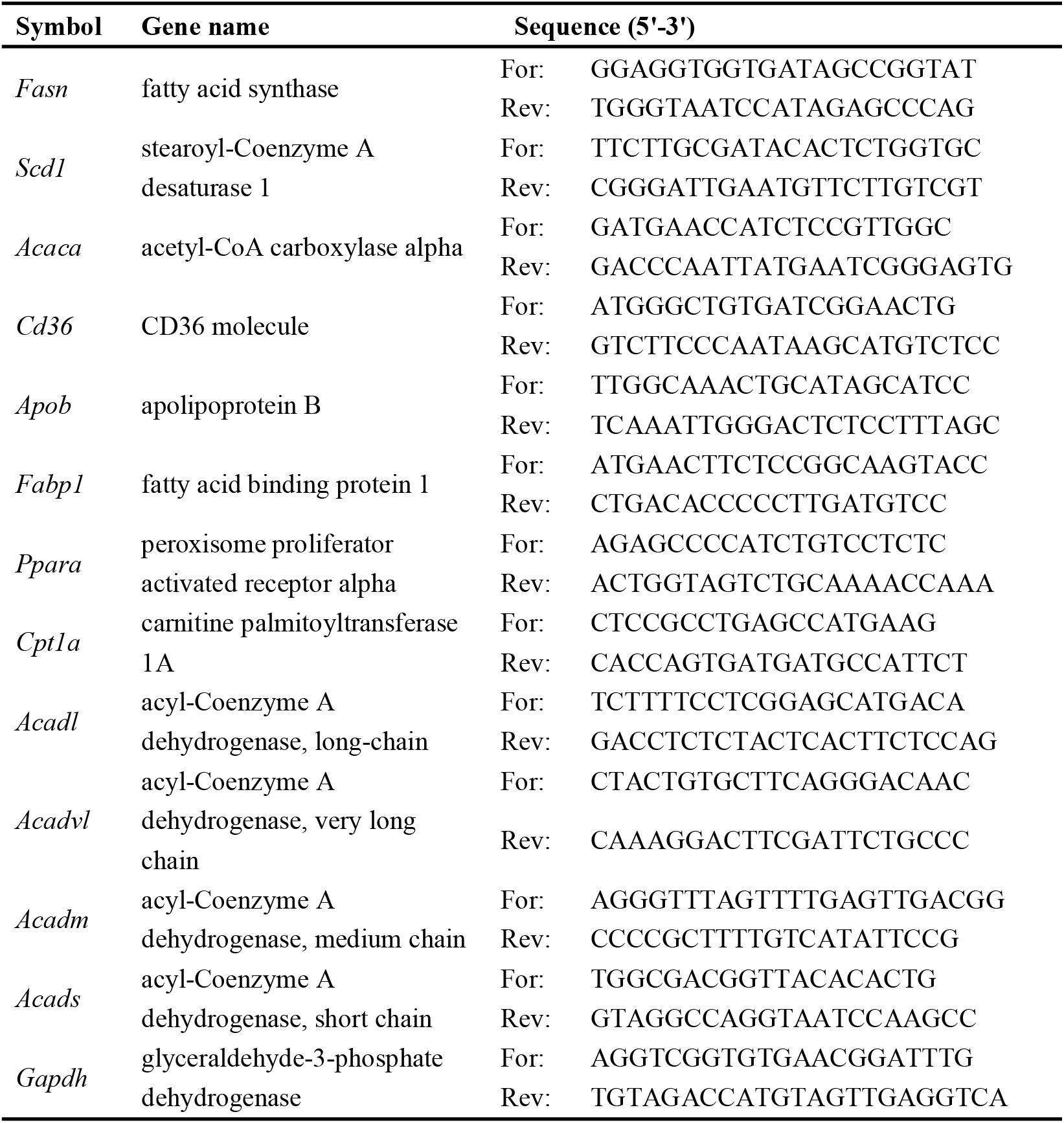
The list of primers used to quantify mRNA.

## Notes

### Competing Interest Statement

The authors have declared no competing interest.

